# Dual blockade of IL-18 and IL-4Rα by a bispecific antibody for improved treatment of atopic dermatitis

**DOI:** 10.64898/2026.10.11.758381

**Authors:** Lijuan Zhang, Zhan Meng, Hui Bo, Ye Tian, Weicai Mou, Xianqiang Ping, Junya Dong, You Leo Li

## Abstract

**Background:** Atopic dermatitis (AD) is a chronic inflammatory dermatosis primarily driven by type 2 immunity, with additional type 1 and type 3 components that may limit the benifit of IL-4Rα blockade alone. Interleukin-18 (IL-18), an epithelial alarmin genetically associated with AD and correlated with disease severity, has been implicated as an upstream amplifier of this mixed inflammatory response.

**Objective:** To assess whether concurrent IL-18 neutralization and IL-4Rα blockade with a bispecific antibody provides broader and more balanced suppression of AD-relevant type 1, type 2, and type 3 pathways than single-target inhibition.

**Methods:** We generated FJ-0125, a bispecific antibody comprising human IL-18 binding protein fused to a novel humanized anti-IL-4Rα antibody, and characterized its binding, signaling, and functional activity in vitro and in vivo. The activity was evaluated in human group 2 innate lymphoid cell and primed Th2 cell assays, a keratinocyte–dendritic cell–T cell coculture system, and an MC903/Staphylococcus aureus–induced dermatitis model in human IL-4/IL-4Rα knock-in mice.

**Results:** FJ-0125 bound to IL-18 and IL-4Rα with high affinity, engaged both targets, inhibited IL-4/IL-13 signaling with dupilumab-comparable potency, and neutralized human and mouse IL-18. In vitro, FJ-0125 more effectively suppressed IL-18–driven lymphocyte activation and type 1, type 2, and type 3 cytokine production than IL-18BP-Fc or IL-4Rα monotherapy. In vivo, FJ-0125 reduced multitype cytokines, improved dermatitis, lowered serum IgE, and abrogated the paradoxical IL-22 increase observed with selective IL-4Rα blockade.

**Conclusion:** Dual IL-18/IL-4Rα blockade with FJ-0125 achieves coordinated suppression of multitype inflammation and represents a mechanistically substantiated therapeutic strategy for AD.

**Highlights:** FJ-0125 is a bispecific antibody fusing IL-18 binding protein to anti–IL-4Rα.

FJ-0125 neutralizes IL-18 and blocks IL-4/IL-13 signaling with high potency.

Dual blockade broadly suppresses type 1, 2, and 3 inflammation in vitro.

FJ-0125 outperforms dupilumab in an MC903/S. aureus dermatitis model.

FJ-0125 prevents the IL-22 rebound seen with selective IL-4Rα blockade.

## Introduction

Atopic dermatitis (AD) is a chronic, relapsing inflammatory skin disease that affects approximately 2–5% of adults and a substantial proportion of children worldwide, imposing a major burden on quality of life [1]. Although traditionally regarded as a type 2–dominant disorder driven by interleukin-4 (IL-4) and IL-13, AD is now recognized as a heterogeneous, multitype inflammatory disease in which type 1 (IFN-γ) and type 3 (IL-17/IL-22) responses coexist with type 2 pathways and vary according to disease stage, anatomical site, age, and ethnicity [2–4]. A defective epidermal barrier—exemplified by filaggrin loss-of-function variants—and cutaneous dysbiosis characterized by Staphylococcus aureus colonization further shape this inflammatory milieu [5,6]. This immunological heterogeneity is increasingly considered a key determinant of the variable and often incomplete clinical responses observed with therapies that target a single cytokine pathway.

The clinical success of dupilumab, a monoclonal antibody that blocks the shared IL-4 receptor α-chain (IL-4Rα) and thereby inhibits both IL-4 and IL-13 signaling, has validated the type 2 axis as a central therapeutic target in AD [7]. However, a considerable proportion of patients achieve only partial responses, experience loss of efficacy over time, or develop paradoxical inflammatory manifestations, and residual disease activity is frequently associated with persistent non–type 2 inflammation [2,8]. A growing number of approved and late-stage agents—including the anti–IL-13 antibodies tralokinumab and lebrikizumab, the anti–IL-31 receptor antibody nemolizumab, the OX40/OX40L inhibitors rocatinlimab and amlitelimab, and Janus kinase inhibitors such as upadacitinib—each modulate a single dominant pathway and thus face similar constraints [9–14]. These limitations have prompted efforts to identify complementary upstream drivers that sustain the broader inflammatory network and can be co-targeted to deepen and broaden clinical responses.

Interleukin-18 (IL-18), an IL-1 family alarmin, has emerged as one such candidate [15]. IL-18 is constitutively synthesized by keratinocytes as an inactive precursor and is released in its mature form upon inflammasome activation triggered by barrier disruption and microbial cues, particularly S. aureus, which colonizes the majority of AD lesions [6,16–19]. Multiple lines of human evidence implicate IL-18 in AD pathogenesis: genetic variants near IL18R1 are associated with AD risk [20]; Mendelian randomization analyses support a causal role for IL-18 in eczema [21,22]; and IL-18 protein levels in serum and skin correlate with disease severity [15]. Mechanistically, IL-18 activates group 2 innate lymphoid cells (ILC2s) and basophils and, through IL-9–mediated upregulation of its receptor, licenses pathogenic Th2 cells to secrete IL-13, IL-5, and IL-22 [23–26]. The therapeutic relevance of this pathway is supported by recent clinical proof-of-concept for the anti–IL-18 antibody aletekitug in moderate-to-severe AD [27].

Despite these convergent data, IL-18 and the IL-4/IL-13 axis have largely been pursued as independent therapeutic targets. Several observations suggest that these pathways are mechanistically intertwined and may be more effectively controlled in combination. IL-4, acting via STAT6, is a critical negative regulator of IL-18Rα expression on activated CD4⁺ T cells [28]; conversely, IL-4Rα blockade has been associated with compensatory increases in IL-18 and with persistent or emergent IL-22/type 3 inflammation in patients [29–31]. Baseline IL-18 and IL-22 levels have further been linked to dupilumab treatment outcomes [32]. Together, these findings raise the possibility that selective IL-4Rα blockade, while suppressing type 2 inflammation, may leave an IL-18–driven type 1/type 3 program insufficiently controlled or even accentuated.

Therefore, we hypothesized that simultaneous neutralization of IL-18 and blockade of IL-4Rα within a single molecule could achieve broader and more balanced suppression of the multitype inflammation that characterizes AD. Here, we describe FJ-0125, a bispecific antibody generated by fusing human IL-18 binding protein (IL18BP) to a novel humanized anti–IL-4Rα antibody. We demonstrate that FJ-0125 engages both targets with high affinity, neutralizes both human and mouse IL-18, and, across innate, adaptive, and integrated epithelial–immune in vitro systems, as well as an in vivo model of S. aureus–associated AD, achieves uniform suppression of type 1, type 2, and type 3 inflammation that exceeds that of either single-pathway agent, including abrogation of the IL-22 increase associated with isolated IL-4Rα blockade.

## Materials and Methods

### Materials and Reagents

#### Generation of a humanized anti–IL-4Rα antibody

A novel anti–human IL-4Rα antibody was generated in-house. Mice were immunized with recombinant human IL-4Rα, and hybridomas were produced and screened for high-affinity binding to human IL-4Rα and for blockade of IL-4/IL-13–induced signaling. A lead murine clone was humanized by grafting the complementarity-determining regions onto human variable-region frameworks, with selective framework back-mutations introduced to preserve binding affinity. The humanized anti–IL-4Rα antibody was used as the IL-4Rα–targeting arm for construction of the bispecific antibody. Its binding affinity for human IL-4Rα is reported in Supplementary Figure S1.

#### Antibodies and recombinant proteins

The IL18BP–anti-IL-4Rα bispecific antibody (FJ-0125) was generated in-house as described below. Dupilumab and IL-18BP-Fc were used as benchmark molecules, and isotype-matched human IgG4 was used as a negative control. Recombinant human IL-18 was purchased from Novoprotein (Cat. No. GMP-CH29), and recombinant mouse IL-18 was purchased from Sino Biological (Beijing, China). Recombinant human IL-4Rα-His was obtained from Sino Biological (Cat. No. 10402-H08H-B). Additional recombinant cytokines and proteins used for cell stimulation and binding assays are listed, together with their sources and catalog numbers, in Supplementary Table S1.

#### Bacterial strains

Two Staphylococcus aureus strains were used in this study. For in vitro keratinocyte stimulation, a methicillin-susceptible S. aureus reference strain (ATCC 29213) was used. For the induction of atopic dermatitis (AD)-like inflammation in vivo, a methicillin-resistant S. aureus strain Mu50 (ATCC 700699) was used. Both strains were obtained from the China Center of Industrial Culture Collection (Beijing, China). Bacteria were streaked on Luria–Bertani (LB) agar plates, and a single colony was inoculated into LB broth and cultured at 37 °C with shaking at 200 rpm until the logarithmic growth phase. The bacteria were harvested by centrifugation, washed with sterile phosphate-buffered saline (PBS), and resuspended in PBS. Bacterial concentrations were determined by measuring the optical density at 600 nm (OD₆₀₀) and confirmed by plating serial dilutions on LB agar to enumerate colony-forming units (CFU). For in vitro stimulation, keratinocytes (8 × 10⁴ cells per well) were exposed to S. aureus ATCC 29213 at 1 × 10⁷ CFU per well, corresponding to an infection multiplicity (MOI) of approximately 125:1, for 3 h. The dosing regimen and inoculation of Mu50 used for in vivo induction of AD are described in the Animals and Induction of MC903/Mu50 AD-like dermatitis sections.

### Bispecific Antibody Generation and Characterization

#### Bispecific antibody design and production

The IL18BP–anti–IL-4Rα bispecific antibody (FJ-0125) was generated by combining human IL-18 binding protein (IL18BP) with the humanized anti–IL-4Rα antibody described above, using the knob-into-hole strategy (T366W / T366S-L368A-Y407V) to promote correct heterodimerization. The molecule was built on a human IgG4 backbone carrying S228P, F234A, and L235A mutations to stabilize the hinge and abrogate Fcγ receptor–mediated effector functions. IL-18BP-Fc and isotype control antibodies were constructed in parallel using the same IgG4 backbone. Dupilumab was used as a clinical benchmark comparator. FJ-0125 and control molecules were expressed by transient transfection of ExpiCHO cells according to the manufacturer’s instructions, and culture supernatants were harvested 6 days after transfection. Antibodies were purified by Protein A affinity chromatography followed by preparative size-exclusion chromatography (SEC). Protein purity was confirmed to be >95% prior to use.

#### Binding kinetics by biolayer interferometry (BLI)

The binding kinetics of FJ-0125 to its target antigens were determined by biolayer interferometry using an Octet RED system (Sartorius/ForteBio). Biotinylated FJ-0125 was immobilized onto streptavidin (SA) biosensors, which were then equilibrated in a kinetics buffer (PBS containing 0.02% Tween-20 and 0.1% BSA). Association was measured by incubating the loaded sensors with serial dilutions of recombinant human IL-18 (Novoprotein, Cat. No. GMP-CH29) or recombinant human IL-4Rα-His (Sino Biological; Cat. No. 10402-H08H-B) as the analyte, followed by a dissociation step of approximately 1,400–2,400 s in the kinetics buffer. Reference signals were subtracted, and data were globally fitted using a 1:1 binding model with Octet Analysis Studio version 12.2 to calculate the association rate constant (kₐ), dissociation rate constant (k_d_), and equilibrium dissociation constant (KD).

#### Simultaneous target-binding assay

Concurrent engagement of IL-18 and IL-4Rα by FJ-0125 was assessed by sequential BLI. Biotinylated FJ-0125 was immobilized onto SA biosensors and saturated with the first antigen (recombinant human IL-4Rα-His or human IL-18); the antigen-bound sensors were then transferred into a solution containing the second antigen, and the additional binding response was recorded to determine whether both antigens could be engaged concurrently by the same molecule. Reciprocal assays were performed by reversing the order of antigen addition.

### In Vitro Functional and Neutralization Assays

#### IL-4/IL-13 reporter assay

The blocking activity of FJ-0125 against IL-4Rα-mediated signaling was assessed using an IL-4/IL-13 dual effector reporter cell line (Acro Biosystems, Cat. No. CHEK-ATF075), in which activation of IL-4Rα signaling drives expression of a luciferase reporter gene. Reporter cells were seeded into 96-well plates at 5 × 10⁴ cells/well and stimulated with recombinant human IL-4 (Acro Biosystems, Cat. No. IL4-H4218; final concentration 0.002 μg/mL) or recombinant human IL-13 (Acro Biosystems, Cat. No. IL3-H52H4; final concentration 0.1 μg/mL) in the presence of serial dilutions of FJ-0125, the clinical benchmark antibody dupilumab, IL-18BP-Fc, or an isotype control. After 6 h of incubation, luciferase activity was measured and luminescence was recorded on a microplate reader. Dose–response curves were fitted by nonlinear regression, and IC₅₀ values were determined using a four-parameter logistic model.

#### IL-18 neutralization assay (human PBMCs and mouse splenocytes)

The neutralizing activity of FJ-0125 against IL-18 was evaluated by measuring IFN-γ production in human PBMCs and mouse splenocytes. Human PBMCs (MileCell Bio) were seeded at 1 × 10⁵ cells/well, and mouse splenocytes prepared from human IL-4/human IL-4Rα knock-in mice (C57BL/6 background; Biocytogen) were seeded at 2 × 10⁵ cells/well in 96-well plates. Human PBMCs were stimulated with recombinant human IL-18 (Novoprotein, Cat. No. GMP-CH29; 50 ng/mL) together with recombinant human IL-12 (Sino Biological; 10 ng/mL). For mouse splenocyte assays, two forms of mouse IL-18 were used: wild-type mouse IL-18 (wt IL-18; Sino Biological), which binds both IL-18 receptor 1 (IL-18R1) and IL-18 binding protein (IL-18BP), and a decoy-resistant mouse IL-18 variant (DR-18), which retains equivalent binding to IL-18R1 and activation of IL-18 signaling but does not bind IL-18BP, as previously described by Zhou et al. [33]. DR-18 was generated in-house. Mouse splenocytes were stimulated with wt IL-18 or DR-18 (50 ng/mL) together with recombinant mouse IL-12 (Sino Biological; 10 ng/mL). In both assays, cells were incubated with serial dilutions of FJ-0125, dupilumab, IL-18BP-Fc, or an isotype control. After 48 h, culture supernatants were collected and IFN-γ concentrations were quantified by ELISA (Dakewe Biotech). Dose–response curves were fitted, and IC₅₀ values were calculated using a four-parameter logistic model.

### Keratinocyte and ILC2 Assays

#### Keratinocyte culture and stimulation

Primary normal human epidermal keratinocytes (NHEKs) were cultured in EpiLife medium supplemented with 0.1 ng/mL epidermal growth factor, 10 μg/mL insulin, 0.5 μg/mL hydrocortisone, 50 μg/mL gentamicin, 50 ng/mL amphotericin B, and 0.4% (vol/vol) bovine brain pituitary extract, at 37 °C in a humidified 5% CO₂ incubator. For stimulation, NHEKs (8 × 10⁴ cells per well) were exposed to S. aureus ATCC 29213 at 1 × 10⁷ CFU per well (MOI ≈ 125:1) for 3 h.

#### Preparation of S. aureus-conditioned keratinocyte medium

Following the 3-h stimulation, NHEKs were washed to remove S. aureus and further incubated in fresh medium containing antibiotics for 48 h. The S. aureus-conditioned keratinocyte medium, containing epithelial-derived mediators including TSLP and IL-18, was collected, clarified by centrifugation, and passed through a 0.22-μm filter to remove cellular debris and any residual bacteria. TSLP and IL-18 concentrations in the conditioned medium were quantified by ELISA.

#### Human ILC2 isolation, expansion, and culture

Human group 2 innate lymphoid cells (ILC2s) were isolated from human PBMCs using the EasySep isolation kit (STEMCELL Technologies) according to the manufacturer’s instructions. ILC2s were defined as lineage-negative (Lin⁻) CD127⁺ CRTH2⁺ cells and were subsequently expanded in vitro, as previously described [15, 23]. ILC2s were maintained in RPMI-1640 medium supplemented with 10% FBS, recombinant human IL-2 (10 ng/mL), and recombinant human IL-7 (10 ng/mL).

#### ILC2 activation assay

ILC2s were seeded into 96-well plates at 5 × 10⁴ cells/well and stimulated with recombinant human TSLP (10 ng/mL), IL-18 (10 ng/mL), or their combination, in the presence or absence of FJ-0125 or control molecules. After 48 h, culture supernatants were collected and IL-13 production was quantified by ELISA as a marker of ILC2 activation.

#### Keratinocyte-conditioned medium–ILC2 stimulation assay

Human ILC2s were seeded into 96-well plates at 5 × 104 cells/well and stimulated with *S. aureus*-conditioned keratinocyte medium diluted 1:1 with fresh ILC2 medium in the presence of an isotype control, dupilumab, IL-18BP, or FJ-0125. After 48 h, the culture supernatants were collected and IL-13 production was quantified by ELISA.

### In Vitro Th2 Cell Differentiation and IL-18 Stimulation

#### In vitro Th2 cell differentiation

Naïve CD4⁺ T cells were isolated from healthy donor PBMCs using negative selection (Naïve CD4⁺ T Cell Isolation Kit II; Miltenyi Biotec), yielding a purity of ≥95% CD4⁺CD45RA⁺ cells. Cells were activated with anti-CD3/CD28 Dynabeads (Gibco) at a bead-to-cell ratio of 1:1 and cultured at 1 × 10⁶ cells/mL in complete RPMI-1640 medium supplemented with recombinant human IL-4 (20 ng/mL; PeproTech) and an anti-IFN-γ neutralizing antibody (10 μg/mL; R&D Systems) under Th2-polarizing conditions. Recombinant human IL-2 (20 U/mL; PeproTech) was added on day 2. Cells were expanded for 7 days with medium replenishment every 2–3 days, and the Dynabeads were removed on day 5 using a magnetic separator. Th2 polarization was confirmed by flow cytometry based on CRTH2 and CCR8 expression.

#### IL-18 stimulation and antibody treatment

On day 7, primed Th2 cells were harvested, washed, and reseeded at 2 × 10⁵ cells/well in 96-well U-bottom plates in fresh complete medium without exogenous IL-4 or anti-IFN-γ. Cells were stimulated with recombinant human IL-18 (50 ng/mL; Novoprotein, Cat. No. GMP-CH29) in the presence or absence of the following treatments: (1) the FJ-0125 bispecific antibody (50 nM); (2) recombinant human IL-18BP alone (equimolar); (3) dupilumab (equimolar); or (4) an isotype control antibody. All treatments were added simultaneously with IL-18 at the start of stimulation. Cell-free culture supernatants were collected at 48 h and analyzed for IL-13, IL-5, IFN-γ, and IL-22 by multiplex immunoassay (Luminex; R&D Systems).

### Allogeneic Lymphocyte Reaction (ALR) Assay

#### Generation of monocyte-derived dendritic cells (moDCs)

PBMCs were isolated from healthy donors by Ficoll (Cytiva) density gradient centrifugation. CD14⁺ monocytes were positively selected using CD14 MicroBeads (Miltenyi Biotec) according to the manufacturer’s instructions (purity ≥90%). Monocytes were cultured at 0.5–1 × 10⁶ cells/mL in RPMI-1640 (Gibco) supplemented with 10% heat-inactivated FBS, 2 mM L-glutamine, 100 U/mL penicillin, 100 μg/mL streptomycin, and 50 μM β-mercaptoethanol, in the presence of recombinant human GM-CSF (100 ng/mL; PeproTech) and IL-4 (50 ng/mL; PeproTech). Cultures were maintained for 5–6 days with a half-medium change and cytokine replenishment on day 3. Differentiation into immature moDCs was confirmed by flow cytometry (CD14⁻CD1a⁺HLA-DR⁺CD11c⁺).

#### DC priming

On day 6, immature moDCs were primed for 24 h with recombinant human TSLP (20 ng/mL; R&D Systems) plus recombinant human IL-18 (50 ng/mL; Novoprotein, Cat. No. GMP-CH29). In selected experiments, conditioned medium from S. aureus-stimulated primary human keratinocytes was used as an alternative priming stimulus. DC activation was confirmed by upregulation of OX40L (CD252), CD80, and CD86 by flow cytometry.

#### Allogeneic T cell co-culture

Naïve CD4⁺ T cells were isolated from an allogeneic healthy donor by negative selection using the Naïve CD4⁺ T Cell Isolation Kit II (Miltenyi Biotec) (purity ≥90% CD4⁺CD45RA⁺). Primed moDCs were co-cultured with allogeneic naïve CD4⁺ T cells at a ratio of 1:5 (2 × 10⁴ moDCs : 1 × 10⁵ T cells per well) in 96-well U-bottom plates in a final volume of 200 μL. Recombinant human IL-18 (50 ng/mL; Novoprotein, Cat. No. GMP-CH29) was maintained in the co-culture medium throughout the assay to model sustained epithelial-derived IL-18 exposure. Co-cultures were incubated for 6 days at 37 °C and 5% CO₂. In selected experiments, conditioned medium from S. aureus-stimulated keratinocytes, containing TSLP and IL-18, was used as the co-culture medium.

#### Treatment conditions

Test articles were present throughout both the DC priming and DC–T cell co-culture periods: (1) the FJ-0125 bispecific antibody; (2) recombinant human IL-18BP alone (equimolar to the bispecific); (3) the anti-IL-4Rα monoclonal antibody dupilumab alone (equimolar to the bispecific); or (4) an isotype control antibody. Each condition was tested in triplicate wells per donor pair, with a minimum of three independent donor pairs. Cell-free supernatants were collected on day 6 of co-culture, and concentrations of IL-13, TARC (CCL17), IFN-γ, and IL-22 were measured by individual ELISA (Dakewe Biotech). For each cytokine, results were normalized to the isotype control group and expressed as a percentage of control (% of control).

### In Vivo AD Model

#### Animals

Homozygous human IL-4/human IL-4Rα knock-in mice on a C57BL/6 background (Biocytogen) were used to enable engagement of the human IL-4Rα–targeting arm of FJ-0125 and dupilumab. Mice 6-8-week-old at study initiation were housed under specific pathogen-free conditions in individually ventilated cages under controlled temperature and humidity with a 12-h light/dark cycle, with standard chow and water provided ad libitum, and were acclimatized for at least 7 days before study initiation. Mice were allocated to experimental groups by stratified randomization based on baseline body weight and ear thickness, and investigators performing clinical, histopathological, and biomarker assessments were blinded to treatment allocation. All procedures were reviewed and approved by the Institutional Animal Care and Use Committee of PharmaLegacy Laboratories and were conducted under specific pathogen-free and biosafety level 2 (BSL2) conditions in accordance with institutional biosafety requirements.

#### Bacterial strain and challenge preparation

*S. aureus* Mu50 (ATCC 700699) was used as the challenge strain in this study. A qualified working cell bank was used throughout the study to minimize passage- and batch-associated variability, and strain identity and the expected antimicrobial resistance phenotype were confirmed before study initiation. The bacterial challenge suspension was standardized to an OD₆₀₀ of 1.0, and viable counts were confirmed retrospectively by quantitative culture (approximately 10⁹–10¹⁰ CFU/mL).

#### Induction of MC903/Mu50 AD-like dermatitis

AD-like skin inflammation was induced by repeated topical co-exposure to MC903 and S. aureus Mu50, adapted from previously reported MC903 and MC903–S. aureus models [37, 38]. MC903 (2 nmol per ear per application) dissolved in approximately 20 μL of ethanol was topically applied to each ear. Thirty minutes after each MC903 application, the same ear surfaces were topically challenged with the Mu50 suspension. MC903 and Mu50 were co-administered on the days indicated in Figure 5A, and the tissues were collected on day 20. Vehicle-controlled mice underwent identical handling and received the corresponding MC903 and bacterial vehicle.

#### Treatment and dosing

Treatment was initiated on day 5 of model induction. FJ-0125 (8 mg/kg), dupilumab (10 mg/kg), IL-18BP-Fc (6 mg/kg), or an isotype control antibody (10 mg/kg) was administered by intravenous injection twice weekly (BIW) until study termination. Because the test articles differed in molecular weight, the mg/kg doses were selected to achieve approximately equimolar exposure across groups. The dosing groups and schedule are shown in Figure 5A.

#### Assessment of ear swelling

Ear thickness was measured at baseline and on the days indicated in Figure 5B using a calibrated digital micrometer immediately before topical treatment on challenge days. Three measurements were obtained from a predefined region of each ear and averaged to generate an ear-level value; the mean value from both ears was used as the animal-level measurement.

#### Histopathological analysis

At the end of the study, ear tissues were collected, fixed in neutral-buffered formalin, paraffin-embedded, sectioned, and stained with hematoxylin and eosin (H&E). Representative sections are shown in the figure legend.

#### Measurement of tissue cytokines and serum IgE

Ear tissues were homogenized in protein-extraction buffer supplemented with protease inhibitors. Concentrations of IL-6, IL-4, IFN-γ, IL-22, IL-13, and IL-17 in tissue homogenates were measured by ELISA (Dakewe Biotech) and normalized to total protein concentration. Serum IgE levels were determined by ELISA (Dakewe Biotech).

### Statistical Analysis

For in vitro assays, data are presented as mean ± SEM from at least three independent experiments or donor pairs. Comparisons were performed using one-way ANOVA with Tukey’s post hoc test or a paired t-test, as appropriate. For the in vivo study, ear thickness on day 17 and day 20 was analyzed separately by one-way ANOVA with Tukey’s multiple-comparison test. Endpoint tissue cytokine and serum IgE data were log-transformed and analyzed by one-way ANOVA with Tukey’s multiple-comparison test. Data are presented as mean ± SEM, and P < 0.05 was considered statistically significant. *P < 0.05; **P < 0.01; ***P < 0.001; ****P < 0.0001. All analyses were performed using GraphPad Prism 10.5.0.

## Results

### Discovery and humanization of a novel anti–IL-4Rα antibody

To generate the IL-4Rα–targeting arm of the bispecific antibody, we immunized mice with recombinant human IL-4Rα and screened hybridomas for high-affinity binding to human IL-4Rα and for blockade of IL-4/IL-13 signaling. A lead murine clone was humanized by grafting the complementarity-determining regions onto human variable-region frameworks, followed by selective framework back-mutations to preserve affinity. The resulting humanized antibody (clone hIL4R-0125) bound recombinant human IL-4Rα with high affinity (KD = 5.0 × 10⁻¹² M; Supplementary Fig. S1) and effectively inhibited IL-4/IL-13–induced signaling. This humanized anti–IL-4Rα antibody (hIL4R-0125) was used as the IL-4Rα–targeting arm of the bispecific molecule.

### Design and dual-target binding properties of the IL18BP–anti–IL-4Rα bispecific antibody FJ-0125

To simultaneously target the IL-18 and IL-4/IL-13 inflammatory axes implicated in AD, we engineered a bispecific antibody, FJ-0125, by fusing human IL-18BP to a humanized anti–IL-4Rα antibody (clone hIL4R-0125) using a knob-into-hole Fc heterodimerization strategy on an effector-silenced human IgG4 backbone (Fig. 1A).

**Figure 1.**
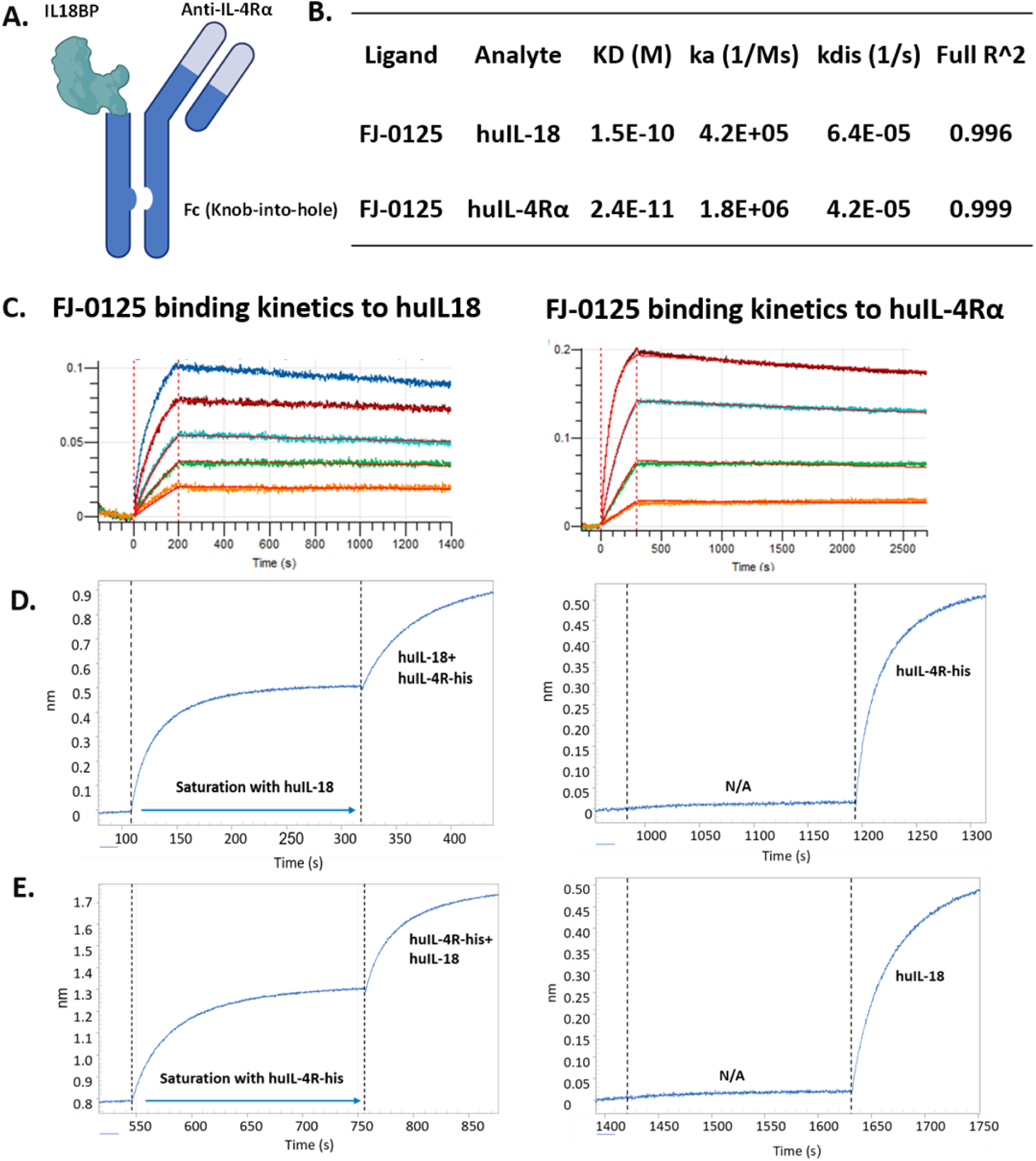
Design and dual-target binding characterization of the IL18BP–anti-IL-4Rα bispecific antibody FJ-0125. (A) Schematic representation of the IL18BP–anti-IL-4Rα bispecific antibody generated using a knob-into-hole Fc heterodimerization strategy on an effector-silenced human IgG4 backbone. (B) Kinetic parameters (ka, kd, KD) of FJ-0125 binding to recombinant human IL-18 and human IL-4Rα, determined by biolayer interferometry (BLI). (C) Representative BLI sensorgrams showing binding of FJ-0125 to recombinant human IL-18 and human IL-4Rα. Sensorgrams were globally fitted using a 1:1 binding model. (D, E) Sequential binding analysis of FJ-0125 with recombinant human IL-18 and human IL-4Rα. Binding responses were recorded following sequential exposure to the two antigens in either order, demonstrating simultaneous dual target engagement.

We characterized the binding properties of FJ-0125 using biolayer interferometry (BLI). FJ-0125 bound recombinant human IL-18 and human IL-4Rα with high affinity, with equilibrium dissociation constants (KD) of 1.5 × 10⁻¹⁰ M and 2.4 × 10⁻¹¹ M, respectively (Fig. 1B, C). To determine whether both targets could be engaged concurrently, we performed sequential binding assays. Following saturation with one antigen, FJ-0125 retained the capacity to bind the second antigen, irrespective of the order of antigen addition (Fig. 1D, E). Thus, fusion of IL18BP to the anti–IL-4Rα scaffold preserves high-affinity, simultaneous recognition of both IL-18 and IL-4Rα by a single molecule.

### FJ-0125 blocks IL-4Rα signaling and neutralizes both human and mouse IL-18

Next, we evaluated the dual functional activity of FJ-0125 in vitro. In an IL-4/IL-13 dual effector reporter cell line, FJ-0125 inhibited both IL-4– and IL-13–induced signaling in a dose-dependent manner, with IC₅₀ values of 0.33 nM and 0.15 nM, respectively, comparable to dupilumab (0.27 nM and 0.10 nM, respectively; Fig. 2A, B). As expected, IL-18BP-Fc, which does not target IL-4Rα, showed no inhibitory effect.

**Figure 2.**
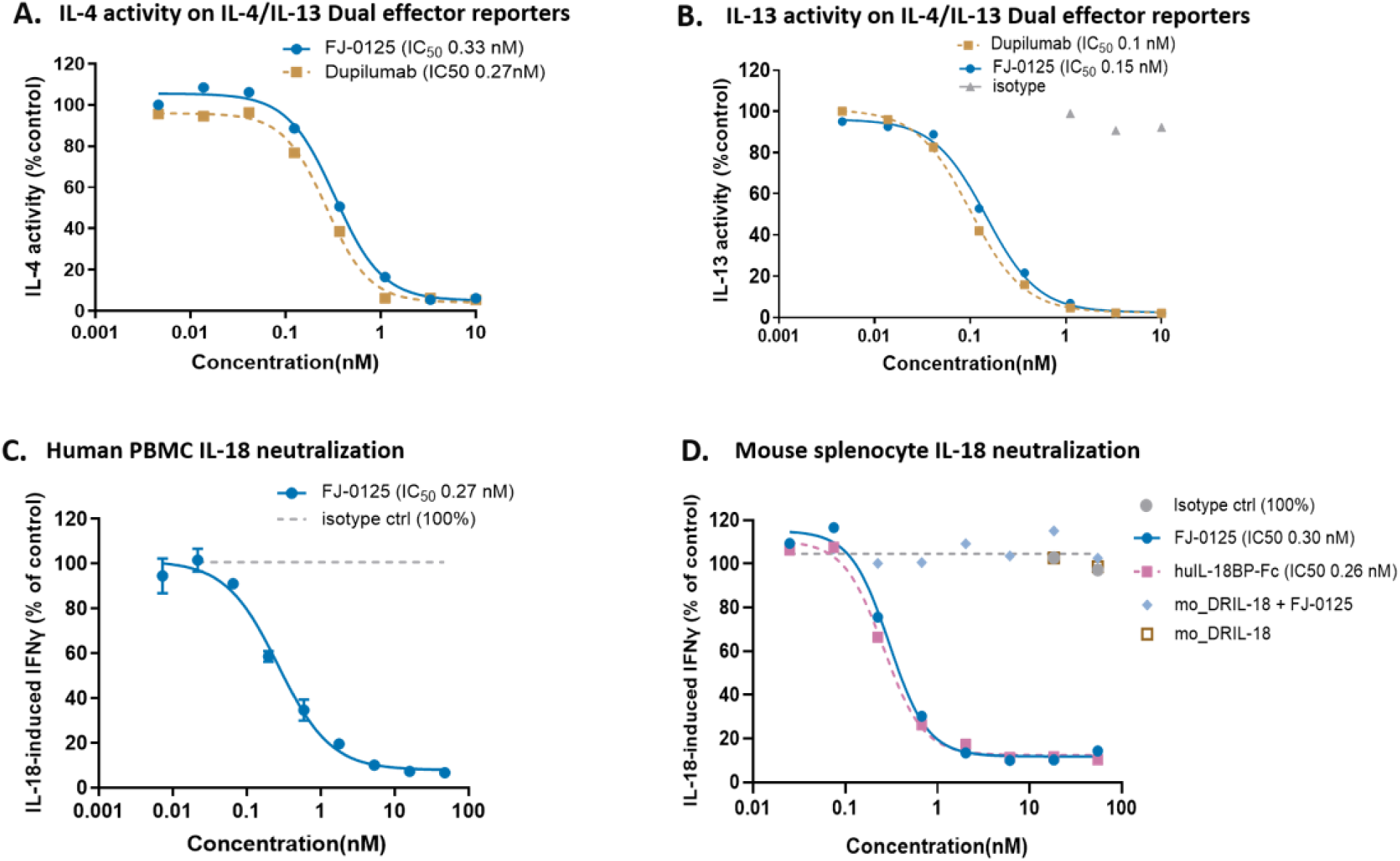
Functional characterization of the IL18BP–anti-IL-4Rα bispecific antibody FJ-0125. (A, B) Inhibition of IL-4– (A) and IL-13–induced (B) signaling by FJ-0125 and the clinical benchmark anti-IL-4Rα antibody dupilumab in an IL-4/IL-13 dual-effector reporter cell assay. (C) Neutralization of human IL-18–induced IFN-γ production by FJ-0125 and the indicated control molecules in human PBMCs stimulated with IL-18 in the presence of IL-12. (D) Neutralization of mouse IL-18 activity by FJ-0125 in mouse splenocytes stimulated with wild-type mouse IL-18 (wt IL-18) or decoy-resistant mouse IL-18 variant (DR-18), together with mouse IL-12. IFN-γ levels in the culture supernatants were quantified using ELISA. DR-18, which does not bind IL-18BP but retains IL-18R1 signaling, was used as a specificity control for IL18BP-mediated neutralization. Dose–response curves were fitted using a four-parameter logistic model.

To assess the IL-18-neutralizing activity, we measured IFN-γ production by human PBMCs and mouse splenocytes stimulated with IL-18 in combination with IL-12. FJ-0125 dose-dependently suppressed IL-18-induced IFN-γ production in both human PBMCs (IC₅₀ = 0.27 nM) and mouse splenocytes (IC₅₀ = 0.30 nM), with a potency comparable to that of human IL-18BP-Fc (IC₅₀ = 0.26 nM), demonstrating that FJ-0125 neutralizes both human and mouse IL-18 with similar efficacy (Fig. 2C, D). This cross-species reactivity enabled subsequent evaluation of FJ-0125 in mouse models of AD. In contrast, dupilumab showed no IL-18-neutralizing activity, and the isotype control had no effect on IL-18-induced IFN-γ production.

To confirm that the IL-18–neutralizing activity of FJ-0125 was mediated specifically through its IL18BP module, we stimulated mouse splenocytes with either wild-type IL-18 (wt IL-18) or a decoy-resistant IL-18 variant (DR-18) that retains IL-18R1 signaling but does not bind IL-18BP [33]. FJ-0125 suppressed IFN-γ production induced by wt IL-18 but was unable to block that induced by DR-18 (Fig. 2D), confirming that its IL-18–neutralizing activity is mediated specifically by the IL18BP moiety.

Together, these data show that FJ-0125 integrates high-affinity IL-4Rα blockade—comparable to dupilumab—with IL18BP-mediated neutralization of both human and mouse IL-18 within a single molecule, whereas each single-target comparator was active against only one pathway.

### FJ-0125 suppresses IL-18– and keratinocyte-driven ILC2 activation

Group 2 innate lymphoid cells (ILC2s) are major producers of type 2 cytokines in AD, and both TSLP and IL-18 have been implicated in their activation [23, 24]. We first stimulated human ILC2s with TSLP, IL-18, or a combination of both and measured IL-13 as a readout of activation. TSLP or IL-18 alone induced only modest IL-13 production, whereas their combination triggered a robust, marked, synergistic increase that greatly exceeded the sum of the individual responses (Fig. 3A). FJ-0125 strongly suppressed TSLP-plus-IL-18–induced IL-13 production, reducing it by approximately 80% relative to isotype control; IL-18BP-Fc produced a slightly weaker reduction (∼68%), whereas dupilumab had essentially no effect (∼6% reduction), consistent with the IL-4Rα–independent nature of this stimulus (Fig. 3A).

**Figure 3.**
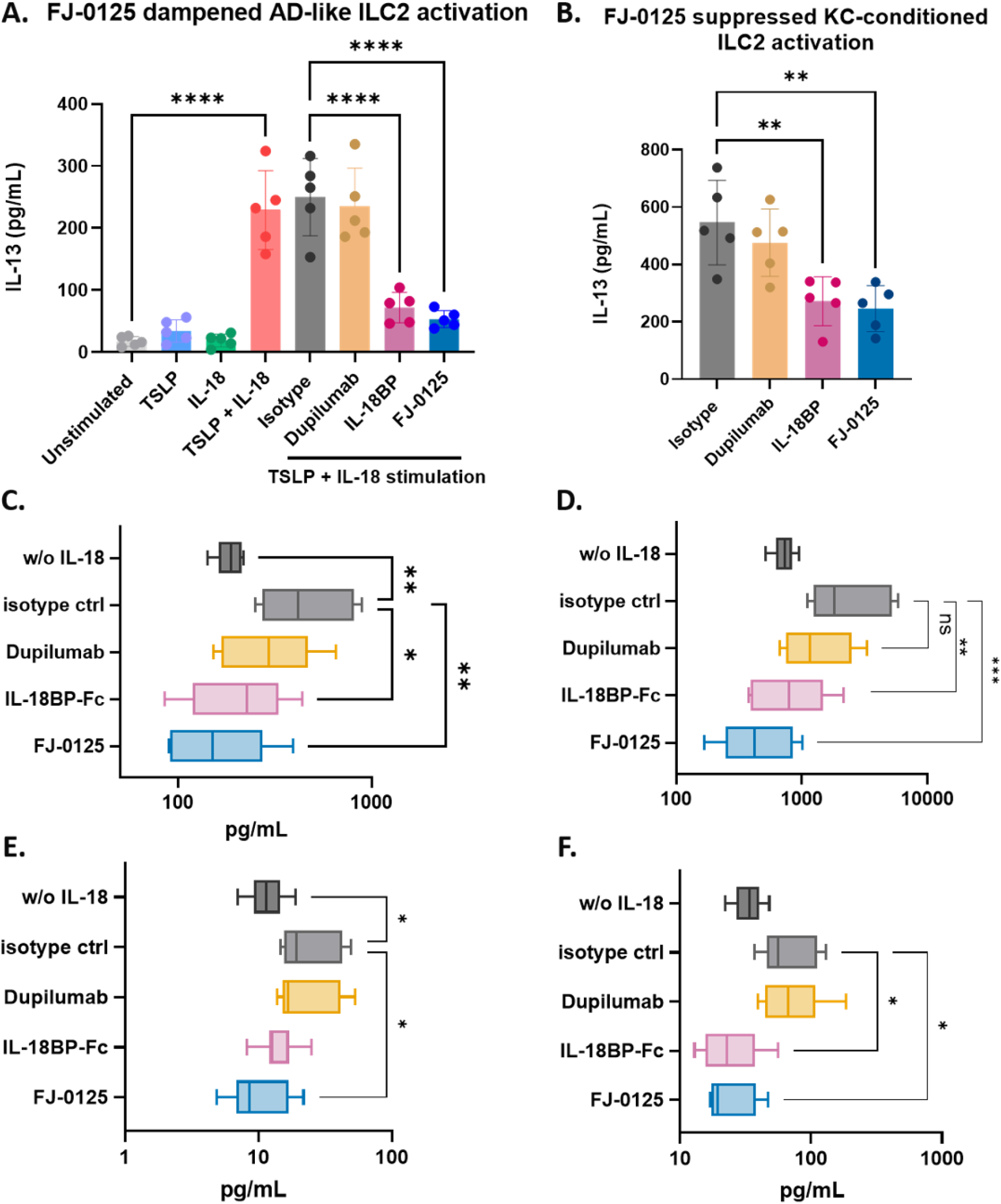
FJ-0125 suppresses IL-18–driven activation of ILC2s and primed Th2 cells. (A) Human ILC2s were stimulated with TSLP, IL-18, or their combination in the presence or absence of FJ-0125 or control molecules. ILC2 activation was evaluated by measuring IL-13 production in the culture supernatant. (B) Human ILC2s were stimulated with S. aureus-conditioned keratinocyte medium in the presence of FJ-0125 or indicated control molecules, and IL-13 production in culture supernatants was measured as an indicator of ILC2 activation. (C–F) Human PBMC-derived Th2 cells were differentiated in vitro under Th2-polarizing conditions and subsequently stimulated with recombinant human IL-18 in the presence of FJ-0125 or indicated control molecules. Cytokine levels in culture supernatants were measured by ELISA. The production of IL-13 (C) and IL-5 (D) was assessed as an indicator of type 2 responses, whereas IFN-γ (E) and IL-22 (F) were measured as markers of type 1- and type 3-associated inflammatory responses, respectively. Data are presented as mean ± SEM. Groups were compared by one-way ANOVA with Tukey’s multiple-comparison test. Detailed statistical analyses are described in the Methods section.

To test FJ-0125 under more pathophysiologically relevant conditions, conditioned medium from S. aureus-stimulated keratinocytes was used as the stimulus. S. aureus stimulation induced keratinocyte secretion of both IL-18 and TSLP (Supplementary Figure S2), identifying keratinocytes as an epithelial source of these ILC2-activating cytokines. Conditioned medium from S. aureus-stimulated keratinocytes induced robust IL-13 production by ILC2s. FJ-0125 significantly suppressed this response (∼55% reduction relative to isotype), as did IL-18BP-Fc (∼49%), whereas dupilumab conferred only minimal inhibition (∼6% reduction; Fig. 3B). Thus, IL-18 and TSLP synergistically drive ILC2 activation, and FJ-0125 inhibits ILC2-derived IL-13 production-primarily through its IL18BP module-under both defined-cytokine and keratinocyte-derived stimulation.

### FJ-0125 suppresses IL-18–driven amplification of multitype cytokine responses in primed Th2 cells

Next, we examined whether FJ-0125 modulated IL-18–driven activation of adaptive type 2 effector cells. Human naïve CD4⁺ T cells were polarized under Th2 conditions for 7 days, and the resulting primed Th2 cells were stimulated with IL-18. Compared with unstimulated cells, IL-18 increased the production of the canonical type 2 cytokines IL-5 and IL-13, as well as the type 1 cytokine IFN-γ and the type 3 cytokine IL-22 (Fig. 3C–F). FJ-0125 significantly reduced all four cytokines relative to isotype control, returning levels to approximately those of unstimulated cells, and IL-18BP-Fc produced a comparable degree of inhibition (Fig. 3C–F). In contrast, dupilumab modestly reduced the type 2 cytokine IL-5 but did not significantly lower IL-13 or IFN-γ, and IL-22 remained at or above isotype-control levels (Fig. 3C–F). Thus, IL-18 amplifies a multitype (type 1/2/3) cytokine program in primed Th2 cells, and FJ-0125—but not IL-4Rα blockade alone—suppresses all three inflammatory axes.

### FJ-0125 achieves balanced suppression of the keratinocyte–dendritic cell–T cell inflammatory axis

After demonstrating that FJ-0125 inhibits IL-18–driven activation of ILC2s and primed Th2 cells, we next assessed its activity in a system integrating the epithelial, dendritic, and T cell compartments. We established an allogeneic lymphocyte reaction (ALR) in which monocyte-derived dendritic cells (moDCs) were primed with conditioned medium from S. aureus–stimulated keratinocytes containing TSLP, IL-18, and other epithelial-derived mediators, and then co-cultured with allogeneic naïve CD4⁺ T cells, thereby recapitulating the epithelial-to-immune signaling cascade implicated in AD (Fig. 4A).

**Figure 4.**
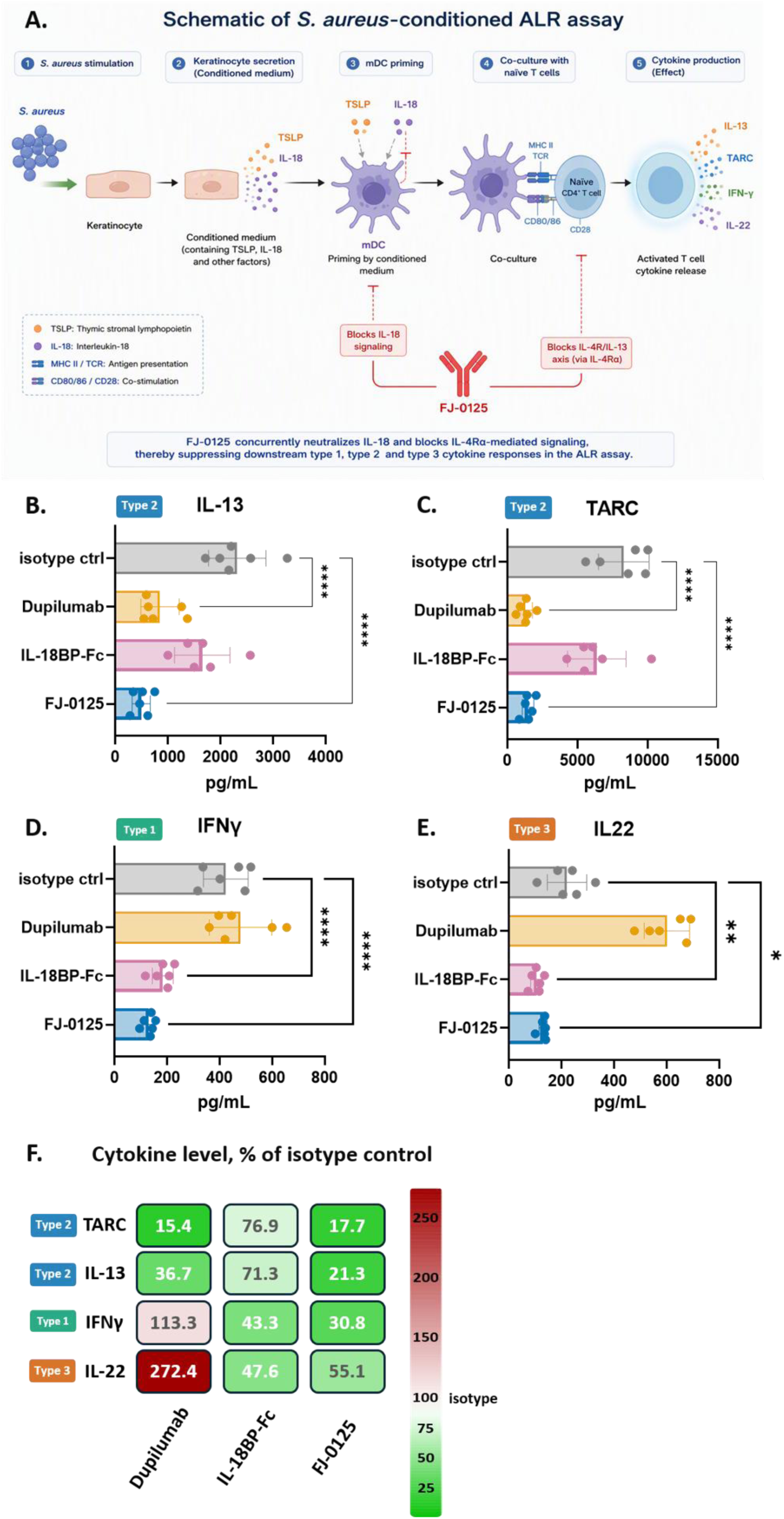
FJ-0125 broadly inhibited keratinocyte-driven dendritic cell–T cell inflammatory responses in an allogeneic lymphocyte reaction (ALR) assay. (A) Schematic representation of the S. aureus-conditioned ALR assay. Human keratinocytes were stimulated with S. aureus to generate conditioned medium containing TSLP, IL-18, and other epithelial-derived mediators. The conditioned medium was used to prime monocyte-derived dendritic cells (moDCs), which were subsequently co-cultured with allogeneic naïve CD4⁺ T cells. Antigen presentation (MHC II/TCR) and co-stimulation (CD80/86–CD28) drive naïve CD4⁺ T cell activation and the production of type 1, type 2, and type 3 cytokines, including IL-13, TARC, IFN-γ, and IL-22. FJ-0125 concurrently neutralizes IL-18 during moDC priming and blocks IL-4Rα-mediated (IL-4/IL-13) signaling during the co-culture phase. Cytokine production in culture supernatants was quantified by ELISA, including IL-13 (B), TARC (C), IFN-γ (D), and IL-22 (E). (F) Heatmap summarizing the cytokine production across the treatment groups. Values represent relative cytokine levels normalized to the isotype control group (set as 100%). Data are presented as mean ± SEM from independent experiments. One-way ANOVA with Tukey ’s multi-comparison test was used for statistical analysis.

**Figure 5.**
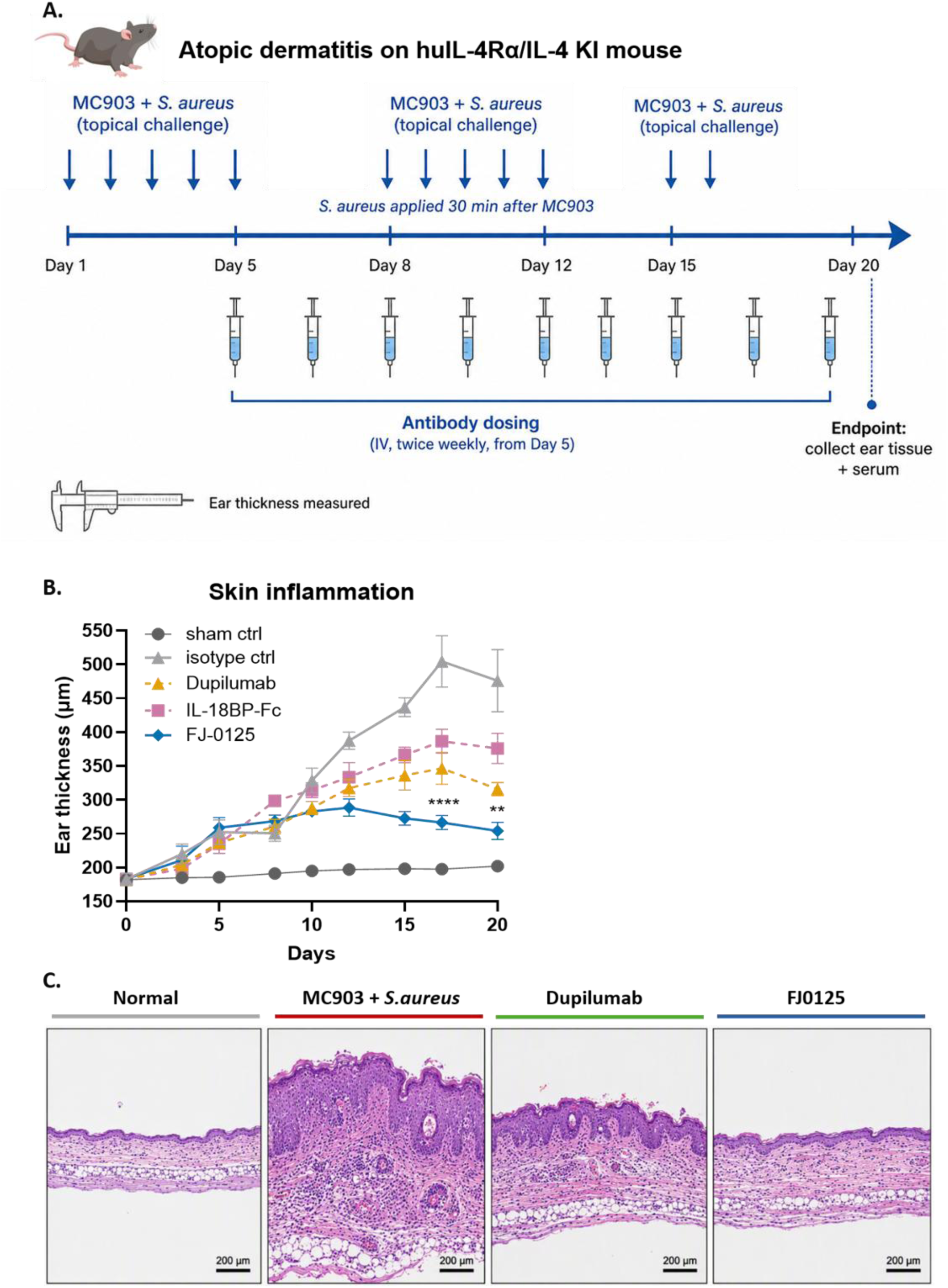
FJ-0125 attenuates ear swelling and skin inflammation in an MC903/S. aureus-induced atopic dermatitis model. (A) Schematic representation of AD model induction and dosing regimen. AD-like skin inflammation was induced in human IL-4/IL-4Rα knock-in mice by topical application of MC903 to the ears, followed 30 min later by topical application of S. aureus at each MC903 administration to mimic epicutaneous bacterial colonization. MC903/S. aureus challenge was applied on the indicated days, and ear tissues were collected at the endpoint (day 20). Beginning on day 5, the mice received FJ-0125, dupilumab, IL-18BP-Fc, or isotype control intravenously twice weekly. (B) Ear thickness measured over the course of the study as an indicator of skin inflammation. Data are shown as mean ± SEM (n = 6 per group). On days 17 and 20, groups were compared by one-way ANOVA with Tukey’s multiple-comparison test; all treatments differed significantly from isotype control (P < 0.001), and stars above bars indicate FJ-0125 versus dupilumab (**P < 0.01, ****P < 0.0001). (C) Representative hematoxylin and eosin (H&E)-stained ear sections at the endpoint from the normal control and MC903/S group. aureus (isotype-treated), dupilumab-treated mice, and FJ-0125-treated mice. Scale bars, 200 μm. Statistical analyses are described in the Methods section.

Activated T cells in this system produce cytokines spanning all three inflammatory axes: IL-13 and TARC (type 2), IFN-γ (type 1), and IL-22 (type 3). FJ-0125 significantly reduced all four cytokines relative to isotype control, lowering them to 17.7% (TARC), 21.3% (IL-13), 30.8% (IFN-γ), and 55.1% (IL-22) of control (Fig. 4B–E). The two single-target comparators each produced an incomplete, skewed profile: dupilumab strongly suppressed the type 2 cytokines (TARC 15.4%, IL-13 36.7% of control) but not IFN-γ (113.3%) and paradoxically increased IL-22 approximately 2.7-fold (272.4%), whereas IL-18BP-Fc reduced the IL-18–dependent cytokines IFN-γ (43.3%) and IL-22 (47.6%) but only modestly affected the type 2 cytokines TARC and IL-13 (76.9% and 71.3% of control) (Fig. 4B–E). The heatmap summary of these responses illustrates the complementary but individually incomplete profiles of the two single-target molecules and the balanced, cross-type suppression observed exclusively with FJ-0125 (Fig. 4F). Notably, the IL-22 increase observed with dupilumab alone was abrogated by FJ-0125.

### FJ-0125 attenuates skin inflammation in a murine model of S. aureus-associated AD

To evaluate the in vivo efficacy of FJ-0125, we used an AD-like model in human IL-4/IL-4Rα knock-in mice, which permits engagement of the human IL-4Rα–targeting arm; the cross-reactivity of the IL18BP module for mouse IL-18 (Fig. 2D) enabled concurrent targeting of both pathways. AD-like dermatitis was induced by repeated topical co-application of MC903 and S. aureus Mu50, and mice were treated with FJ-0125, dupilumab, IL-18BP-Fc, or isotype control (intravenously, twice weekly) beginning on day 5 (Fig. 5A).

Repeated MC903/S. aureus challenge produced progressive ear swelling in isotype-treated mice (with ear thickness increasing from ∼200 μm at baseline to ∼500 μm by day 20), whereas normal control mice showed no appreciable change. FJ-0125 markedly reduced ear thickness compared with isotype at both day 17 and day 20 (P < 0.0001), maintaining ear thickness at ∼250–270 μm—close to that of normal controls (Fig. 5B). In contrast, dupilumab and IL-18BP-Fc conferred only partial protection, with ear thickness remaining significantly higher than in the FJ-0125 group at both time points (P < 0.0001 at day 17 and P < 0.01 at day 20; Fig. 5B).

Histological examination at study termination was consistent with these measurements: isotype-treated mice showed pronounced epidermal hyperplasia and dense dermal inflammatory infiltrate, dupilumab-treated mice showed partial improvement, and FJ-0125–treated mice displayed the greatest reduction in epidermal thickening and dermal inflammation, with an architecture approaching that of normal skin (Fig. 5C).

### FJ-0125 suppresses type 1, type 2, and type 3 cytokines and serum IgE in vivo

To assess whether these clinical and histological improvements were associated with a broad attenuation of the underlying inflammatory milieu, we measured cytokines in ear tissue homogenates and serum IgE at study termination. Consistent with the multitype inflammation of S. aureus–associated AD, isotype-treated mice showed elevated levels of the type 2 cytokines IL-4 and IL-13, the type 1 cytokine IFN-γ, the type 3 cytokines IL-22 and IL-17, and the proinflammatory cytokine IL-6 relative to normal controls (Fig. 6A–F).

**Figure 6.**
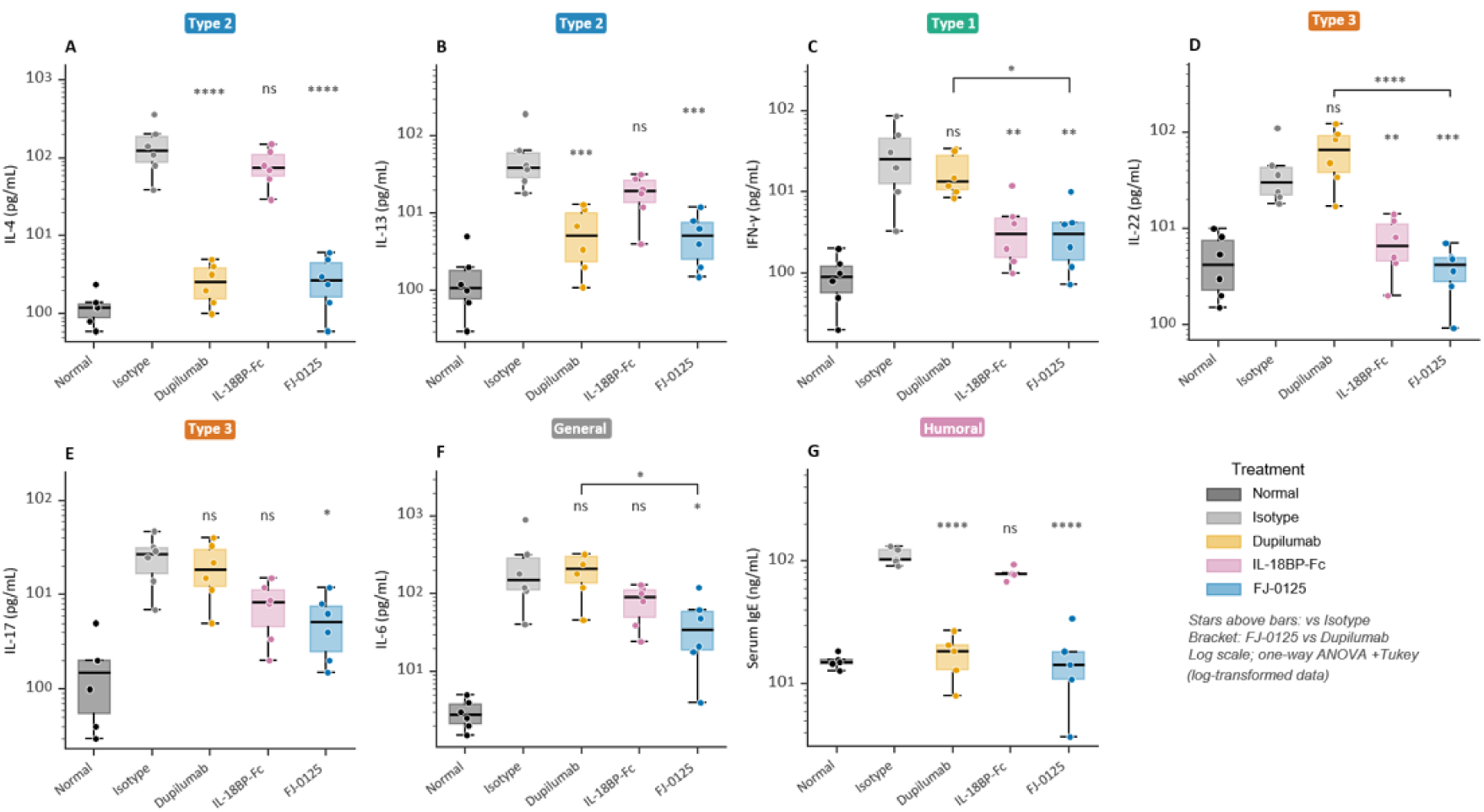
FJ-0125 broadly suppressed type 1, type 2, and type 3 cytokines and serum IgE levels in the MC903/S. aureus-induced atopic dermatitis. (A–F) Cytokine levels in ear tissue homogenates collected at the endpoint from mice in the indicated treatment group. Concentrations of IL-4 (A), IL-13 (B), IFN-γ (C), IL-22 (D), IL-17 (E), and IL-6 (F) were measured by ELISA. IL-4 and IL-13 were assessed as type 2 cytokines, IFN-γ as a type 1 cytokine, and IL-17 and IL-22 as type 3 cytokines; IL-6 was measured as a general inflammatory marker. (G) Serum IgE concentrations at the endpoint, determined by ELISA. Data are presented as box-and-whisker plots with individual data points overlaid (n = 6 per group for tissue cytokines), plotted on a log scale. Groups were compared by one-way ANOVA with Tukey’s multiple-comparison test performed on log-transformed data. Stars above bars indicate significance versus isotype control; brackets indicate FJ-0125 versus dupilumab. *P < 0.05, **P < 0.01, ***P < 0.001, ****P < 0.0001; ns, not significant. Statistical analyses are described in the Methods.

FJ-0125 significantly reduced tissue levels of cytokines across all three inflammatory axes—IL-4 and IL-13 (type 2), IFN-γ (type 1), IL-22 and IL-17 (type 3), and IL-6—compared to the isotype control (Fig. 6A–F). In contrast, the two single-target comparators produced a restricted suppression pattern that mirrored their in vitro profiles. Dupilumab significantly reduced the type 2 cytokines IL-4 and IL-13 but did not significantly lower IFN-γ or IL-6 levels. Tissue IL-22 was not reduced and even trended higher than in isotype-treated mice, although this increase did not reach statistical significance (Fig. 6D). Conversely, IL-18BP-Fc significantly reduced the type 1/type 3 cytokines IFN-γ and IL-22, slightly lowered IL-17 and IL-6, and had little effect on the type 2 cytokines IL-4 and IL-13.

Serum IgE, a systemic biomarker of type 2 activation, was significantly reduced by both FJ-0125 and dupilumab (P < 0.0001) but was not significantly affected by IL-18BP-Fc (Fig. 6G). Thus, only FJ-0125 achieved coordinated suppression of type 1, type 2, and type 3 cytokines together with systemic IgE in vivo.

## Discussion

In this study, we developed FJ-0125, a bispecific antibody that combines a novel humanized anti-IL-4Rα antibody with human IL18BP, and showed that simultaneous blockade of IL-4Rα signaling and neutralization of IL-18 provides broad, balanced suppression of type 1, type 2, and type 3 inflammation in a set of in vitro systems and in an in vivo model of S. aureus–associated AD. Our data support several related conclusions. IL-18 functions as an amplifier of a multitype inflammatory program in AD-relevant effector cells; IL-4Rα blockade and IL-18 neutralization display complementary but individually incomplete activity; and dual blockade by FJ-0125 achieves more uniform cross-type suppression than either single-pathway agent, including abrogation of the paradoxical IL-22 increase associated with isolated IL-4Rα blockade.

Rather than driving a single cytokine axis, IL-18 amplifies a broad inflammatory programme. It synergizes with the epithelial alarmin TSLP to activate ILC2s and drives primed Th2 cells to produce not only the canonical type 2 cytokines IL-5 and IL-13 but also the type 1 cytokine IFN-γ and the type 3 cytokine IL-22. These findings position IL-18, an IL-1 family alarmin released by keratinocytes upon barrier disruption and S. aureus exposure [16–19], as a mechanistic link between epithelial damage, microbial colonization, and the mixed cytokine milieu characteristic of AD. This interpretation aligns with human studies showing that IL-18 activates ILC2s and, through IL-9–mediated upregulation of IL-18R, enables pathogenic Th2 cells to secrete IL-13 and IL-22 [23–26], and with the correlation of IL-18 levels with AD severity [15]. In line with this epithelial–immune relay, S. aureus stimulation induced keratinocyte secretion of both IL-18 and TSLP. Conditioned medium from these cells activated innate (ILC2) and adaptive (dendritic cell–T cell) inflammatory circuits, and both were attenuated by FJ-0125.

The most mechanistically informative finding was the behavior of IL-22 under isolated IL-4Rα blockade. In the integrated keratinocyte–dendritic cell–T cell (ALR) assay, dupilumab not only failed to reduce IL-22 but drove a marked (∼2.7-fold) increase, and a similar upward, though statistically non-significant, trend was observed in vivo. In both settings, concurrent IL-18 neutralization by FJ-0125 abrogated this increase. This observation is concordant with clinical and translational reports that IL-22, and type 3/Th22 inflammation more broadly, can persist or emerge during dupilumab therapy and associated with residual disease activity [30, 32], including a distinct type 22 immune signature in dupilumab-associated head and neck dermatitis [31]. A mechanistic model that links these findings is supported by prior work demonstrating that IL-4 is a key negative regulator of IL-18Rα expression on activated CD4⁺ T cells via STAT6 [28]. By lifting this IL-4/STAT6–dependent repression, IL-4Rα blockade may enhance cellular responsiveness to IL-18, thereby unmasking or amplifying an IL-18–driven type 1/type 3 programme. In line with this hypothesis, dupilumab treatment has been associated with increased local IL-18 production in patients with chronic rhinosinusitis with nasal polyps [29], and baseline IL-18 and IL-22 levels have been linked to dupilumab treatment outcomes in AD [32]. Our data provide a mechanistic rationale for these clinical observations and suggest that concurrent IL-18 neutralization can counteract the type 3 skewing that may accompany selective IL-4Rα blockade.

These findings also situate FJ-0125 within an emerging therapeutic landscape. The clinical proof-of-concept for the anti–IL-18 antibody aletekitug in moderate-to-severe AD [27] validates IL-18 as a therapeutic target, while bispecific approaches that co-target IL-13 with IL-18 [34] or IL-4Rα with IL-31 [35] highlight growing interest in modulating complementary pathways in AD treatment. Most agents now approved or in late-stage development for AD, including inhibitors of IL-13, IL-31R, OX40/OX40L, and Janus kinases [9–14], still act on a single dominant pathway. To our knowledge, FJ-0125 is the first molecule to combine IL-18 neutralization with IL-4Rα blockade in a one entity. This pairing is mechanistically complementary: the anti–IL-4Rα arm restrains the dominant type 2 axis and systemic IgE, whereas the IL18BP arm controls the IL-18–driven type 1/type 3 amplification and the IL-22 rebound that IL-4Rα blockade alone fails to control. The convergence of IL-18 and IL-4/IL-13 signals on shared cellular nodes—particularly dendritic cells and T cells that integrate both inputs—further supports co-targeting the two pathways with a single molecule rather than relying on either agent alone or on a combination of two separate biologics, which would have distinct pharmacokinetics, manufacturing, and dosing considerations [36].

The cross-reactivity of the IL18BP module for both human and mouse IL-18, together with human IL-4/IL-4Rα knock-in mice, allowed both arms of FJ-0125 to be engaged within a single immunocompetent in vivo model. This was important for assessing dual-target efficacy. In the MC903/S. aureus model, which extends the established MC903 model [37] and reproduces the multitype cytokine environment of S. aureus–associated AD [38], FJ-0125 reduced ear swelling and histological inflammation more effectively than either dupilumab or IL-18BP-Fc and achieved uniform suppression of type 1, type 2, and type 3 tissue cytokines together with systemic IgE. The in vivo superiority of FJ-0125 over equimolar single-target comparators supports the view that the improved efficacy derives from simultaneous dual-pathway engagement rather than from either activity alone.

Several limitations merit mention. The epitope of the anti-IL-4Rα arm and a complete developability profile (pharmacokinetics, stability, and immunogenicity) remain to be established, and efficacy was demonstrated in a single MC903/S. aureus knock-in mouse model; confirmation in additional models and longer-term studies will be informative. Given the established role of IL-18 in antimicrobial and antitumor immunity [33,39], the impact of sustained IL-18 neutralization on host defense will also warrant evaluation in longer-term studies, as is standard for immunomodulatory therapies. These data will help define the therapeutic window and clinical potential of dual IL-18/IL-4Rα blockade in AD.

In summary, our findings support IL-18 as a central amplifier of multitype inflammation in S. aureus–associated AD and demonstrate that a bispecific antibody co-targeting IL-18 and IL-4Rα achieves broad, balanced suppression of type 1, type 2, and type 3 inflammation, which is superior to that achieved by single-pathway blockade. FJ-0125 therefore emerges as a mechanistically grounded candidate for AD treatment, with particular promise for patients whose disease is incompletely controlled by IL-4Rα blockade alone.

## Funding

This work was supported in part by Ferry Jam Biotechnology Co., Ltd. Employees of the funder contributed to the production and biophysical characterization of bispecific antibodies, in vivo pharmacology studies, and preparation of the manuscript and are included as authors; their specific contributions are described in the Author Contributions statement.

## Author Contributions

Lijuan Zhang designed the FJ-0125 molecule, conceived and designed the overall study, and drafted the manuscript. Zhan Meng performed the in vitro binding and reporter assays. Hui Bo, Ye Tian, and Junya Dong performed the in vitro functional assays and contributed to data collection and analysis. Xianqiang Ping produced and characterized the bispecific antibody and performed the in vivo pharmacodynamic evaluation. Weicai Mou performed the *Staphylococcus aureus*–related in vitro experiments. You Leo Li contributed to molecular design and to manuscript preparation and editing. All authors read and approved the final manuscript.

## Data Availability

The data supporting the findings of this study are available in the article and its supplementary materials. Raw data are available from the corresponding author upon reasonable requests.

## Ethics and Consent

Human peripheral blood mononuclear cells from healthy donors were obtained from a commercial supplier (MileCell Bio); donor informed consent and ethical review were handled by the supplier.

## Animal Research

All animal procedures were reviewed and approved by the Institutional Animal Care and Use Committee of PharmaLegacy Laboratories and were conducted under specific pathogen-free and biosafety level 2 conditions.

## Sex and Gender Considerations (SGBA)

Sex as a biological variable: Only female mice were used; therefore, the findings may not be generalizable to males. Human PBMCs were obtained from commercial healthy donor sources without sex stratification.

---

**Supplemental Figure S1.**
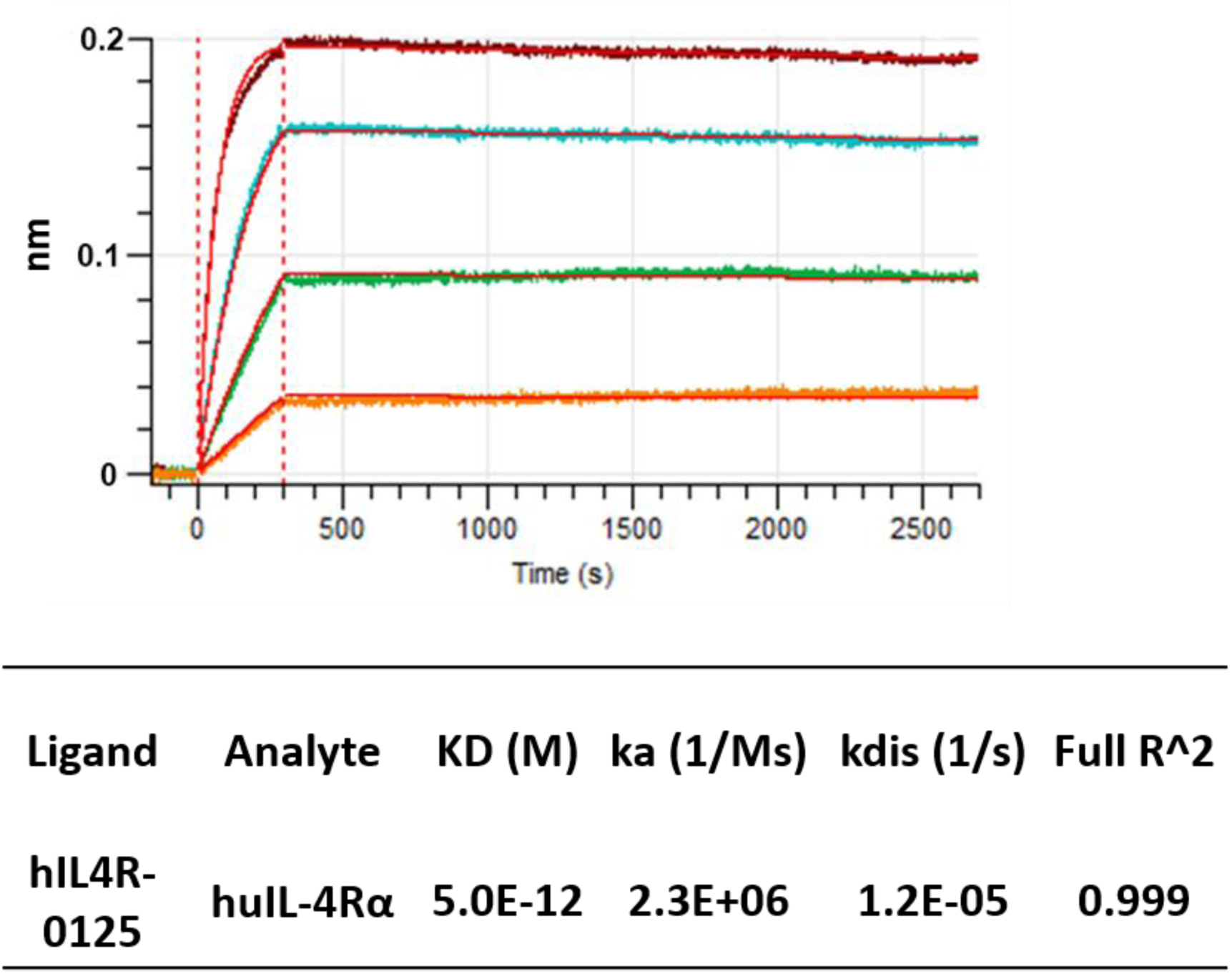

**Supplemental Figure S2.**
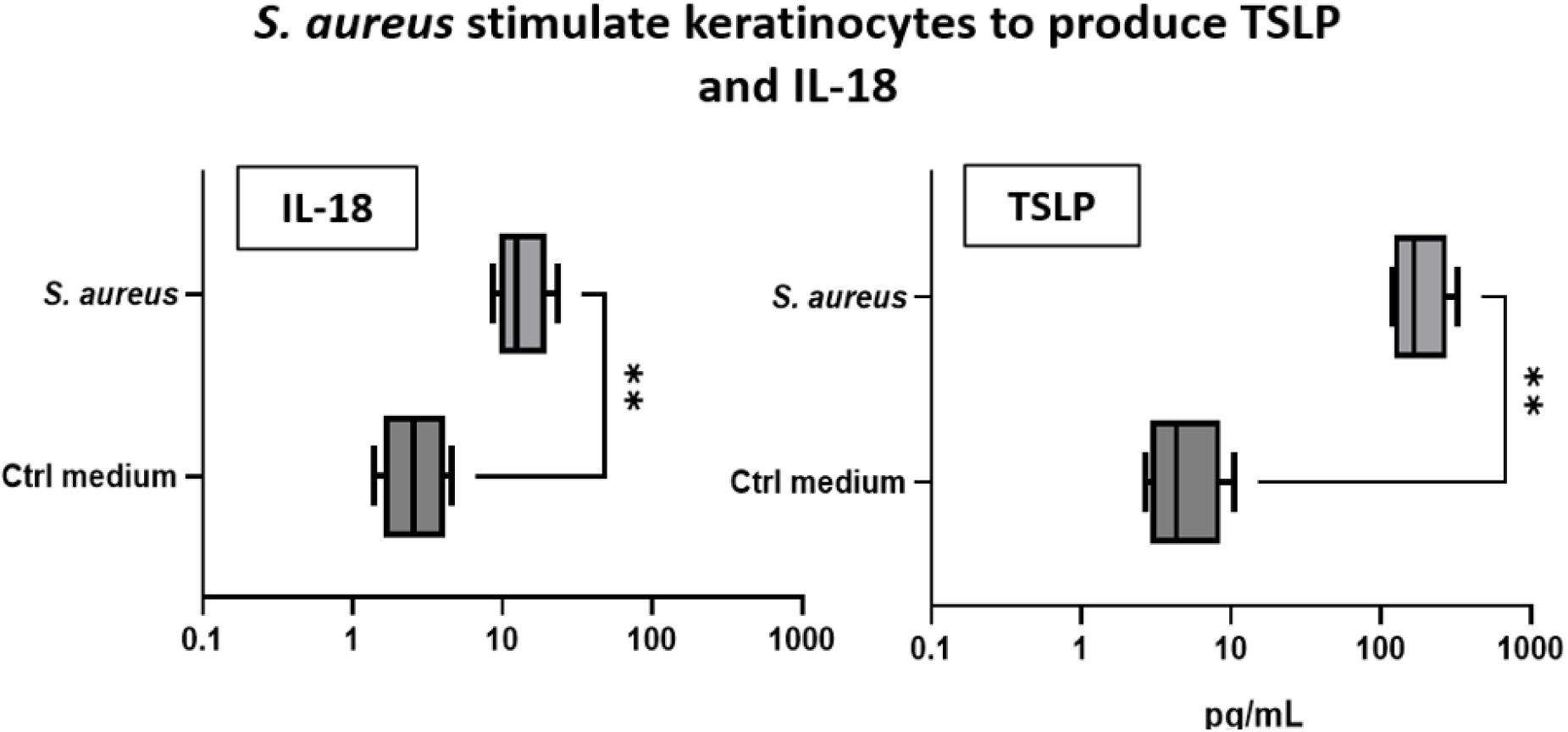
Human keratinocytes were stimulated with S. aureus, and TSLP and IL-18 concentrations in culture supernatants were quantified by ELISA.

**Supplementary Table S1.**
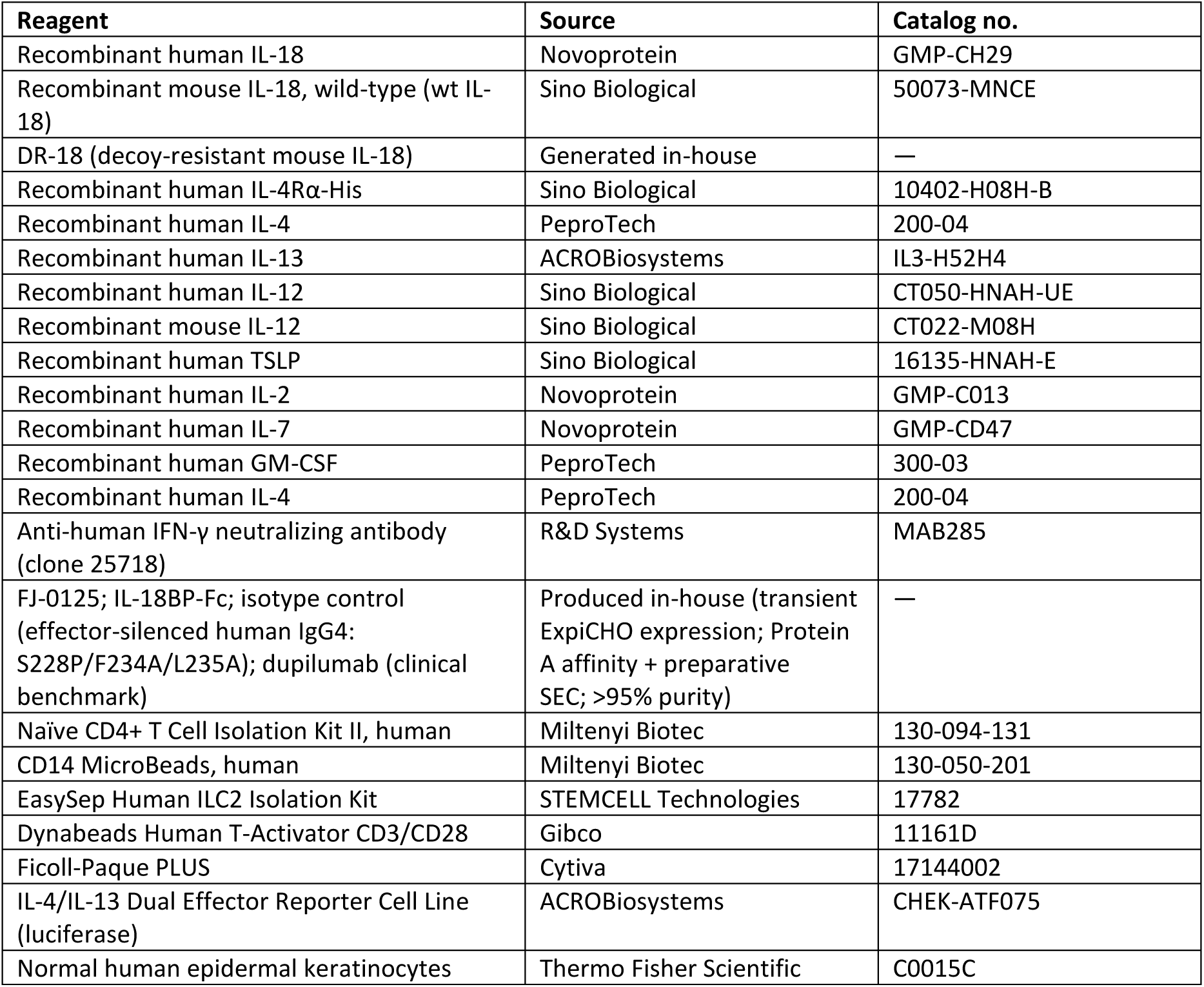
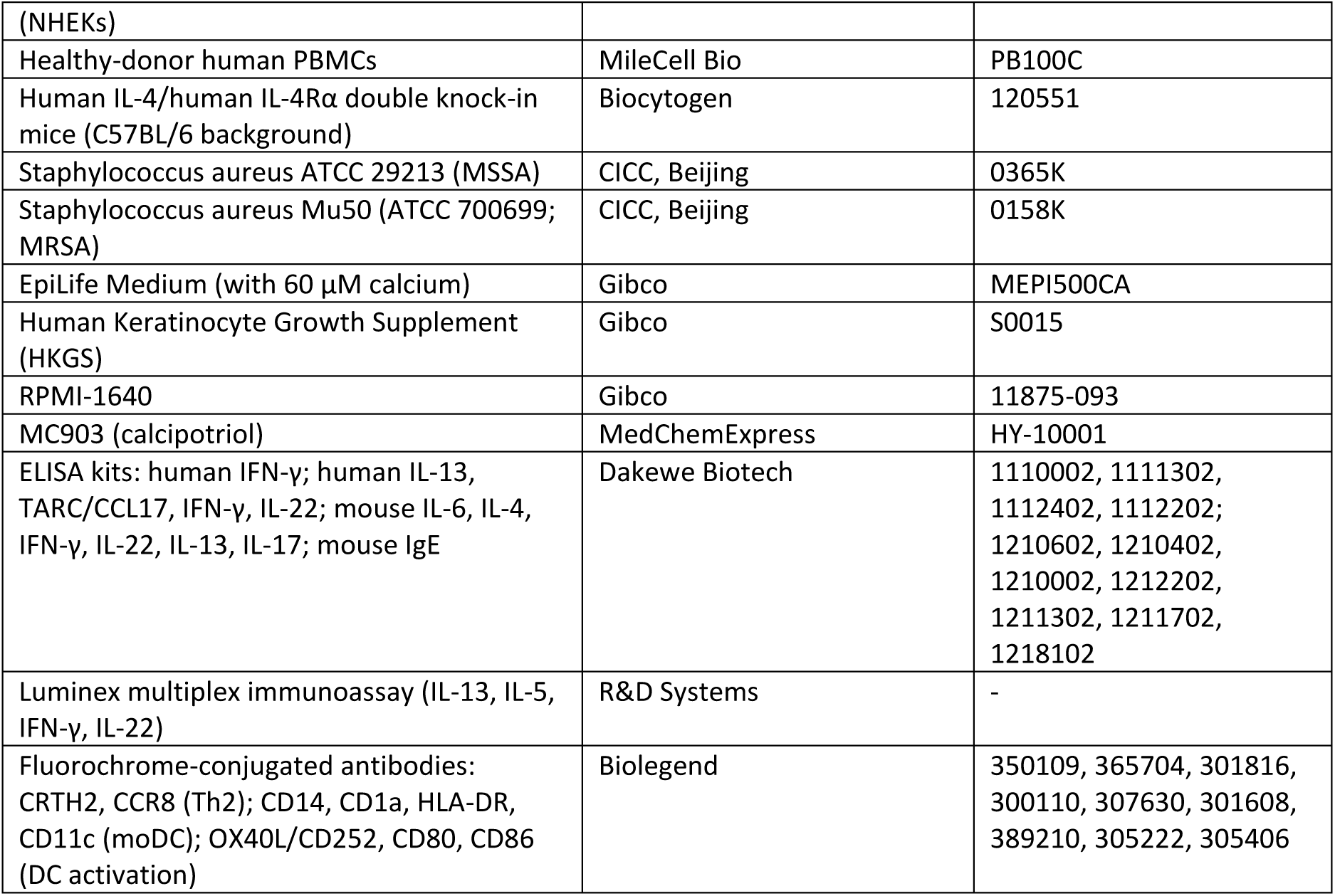
Key reagents, kits, cells, and biological materials.

## Notes

### Competing Interest Statement

X. P. is an employee of Ferry Jam Biotechnology Co., Ltd., which is developing FJ0125. The remaining authors declare no competing interests. Ferry Jam Biotechnology Co., Ltd. has filed a patent application related to FJ0125.

