## supplemental figure 1 and 2 for "Dual blockade of IL-18 and IL-4Rα by a bispecific antibody for improved treatment of atopic dermatitis"

### Supplemental Figure S1

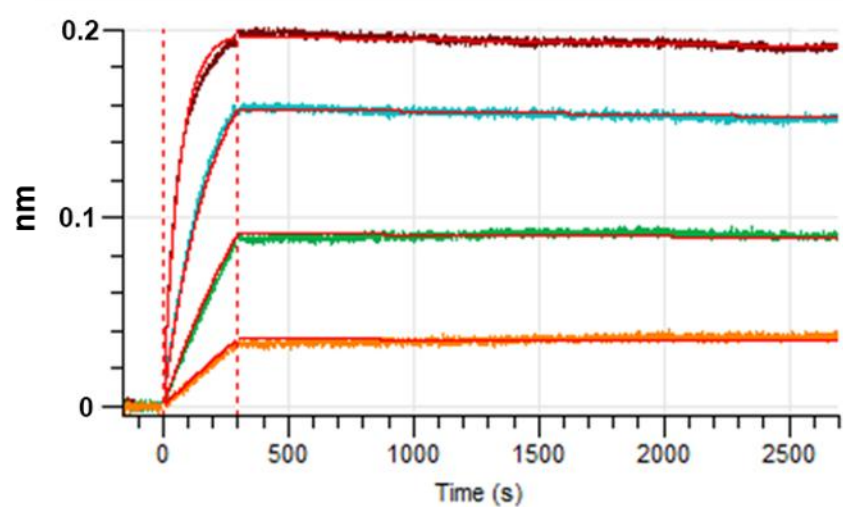

| Ligand | Analyte | KD (M) | ka (1/Ms) | kdis (1/s) | Full R<sup>2</sup> |
| --- | --- | --- | --- | --- | --- |
| hIL4R-0125 | huIL-4Rα | 5.0E-12 | 2.3E+06 | 1.2E-05 | 0.999 |

### Supplemental Figure S2

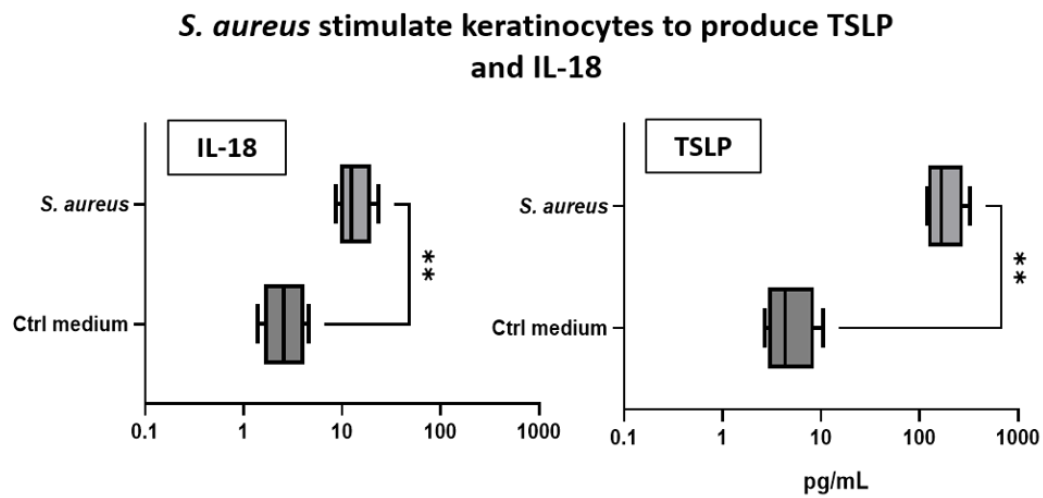

Figure S2. Human keratinocytes were stimulated with *S. aureus*, and TSLP and IL-18 concentrations in culture supernatants were quantified by ELISA.
